# Network hubs in root-associated fungal metacommunities

**DOI:** 10.1101/270371

**Authors:** Hirokazu Toju, Akifumi S. Tanabe, Hirotoshi Sato

## Abstract

**Background:** Although a number of recent studies have uncovered remarkable diversity of microbes associated with plants, understanding and managing dynamics of plant microbiomes remain major scientific challenges. In this respect, network analytical methods have provided a basis for exploring “hub” microbial species, which potentially organize community-scale processes of plant-microbe interactions.

**Methods:** By compiling Illumina sequencing data of root-associated fungi in eight forest ecosystems across the Japanese Archipelago, we explored hubs within “metacommunity-scale” networks of plant-fungus associations. In total, the metadata included 8,080 fungal operational taxonomic units (OTUs) detected from 227 local populations of 150 plant species/taxa.

**Results:** Few fungal OTUs were common across all the eight forests. However, in each metacommunity-scale network representing northern four localities or southern four localities, diverse mycorrhizal, endophytic, and pathogenic fungi were classified as “metacommunity hubs”, which were detected from diverse host plant taxa throughout a climatic region. Specifically, *Mortierella* (Mortierellales), *Cladophialophora* (Chaetothyriales), *Ilyonectria* (Hypocreales), *Pezicula* (Helotiales), and *Cadophora* (incertae sedis) had broad geographic and host ranges across the northern (cool-temperate) region, while *Saitozyma*/*Cryptococcus* (Tremellales/Trichosporonales) and *Mortierella* as well as some arbuscular mycorrhizal fungi were placed at the central positions of the metacommunity-scale network representing warm-temperate and subtropical forests in southern Japan.

**Conclusions:** The network theoretical framework presented in this study will help us explore prospective fungi and bacteria, which have high potentials for agricultural application to diverse plant species within each climatic region. As some of those fungal taxa with broad geographic and host ranges have been known to increase the growth and pathogen resistance of host plants, further studies elucidating their functional roles are awaited.

## Background

Below-ground fungi in the endosphere and rhizosphere are key drivers of terrestrial ecosystem processes [1–4]. Mycorrhizal fungi, for example, are important partners of most land plant species, enhancing nutritional conditions and pathogen resistance of host plants [5–7]. In reward for the essential physiological services, they receive ca. 20% of net photosynthetic products from plants [8, 9]. Recent studies have also indicated that diverse taxonomic groups of endophytic fungi (e.g., endophytic fungi in the ascomycete orders Helotiales and Chaetothyriales) commonly interact with plant roots, providing soil nitrogen/phosphorous to their hosts [10–14], converting organic nitrogen into inorganic forms in the rhizosphere [15], and increasing plants’ resistance to environmental stresses [16–18]. Because of their fundamental roles, below-ground fungi have been considered as prospective sources of ecosystem-level functioning in forest management, agriculture, and ecosystem restoration [17–20]. However, due to the exceptional diversity of below-ground fungi [21–23] and the extraordinary complexity of below-ground plant-fungus interactions [24–26], we are still at an early stage of managing and manipulating plant-associated microbiomes [27–29].

In disentangling complex webs of below-ground plant-fungus associations, network analyses, which have been originally applied to human relations and the World-Wide Web [30, 31], provide crucial insights. By using network analytical tools, we can infer how plant species in a forest, grassland, or farmland are associated with diverse taxonomic and functional groups of fungi [24, 32–34]. Such information of network structure (topology) can be used to identify “hub” species, which are placed at the center of a network depicting multispecies host-symbiont associations [35] (cf. [34, 36, 37]). Those hubs with broad host/symbiont ranges are expected to play key roles by mediating otherwise discrete ecological processes within a community [19, 24]. For example, although arbuscular mycorrhizal and ectomycorrhizal symbioses have been considered to involve distinct sets of plant and fungal lineages [38] (but see [39, 40]), hub endophytic fungi with broad host ranges may mediate indirect interactions between arbuscular mycorrhizal and ectomycorrhizal plant species through below-ground mycelial connections. As information of plant-associated fungal communities is now easily available with high-throughput DNA sequencing technologies [1, 21, 22], finding hub microbial species out of hundreds or thousands of species within a network has become an important basis for understanding and predicting ecosystem-scale phenomena.

Nonetheless, given that fungi can disperse long distances with spores, conidia, and propagules [41–44], information of local-scale networks alone does not provide thorough insights into below-ground plant-fungus interactions in the wild. In other words, no forests, grasslands, and farmlands are free from perturbations caused by fungi immigrating from other localities [45–49]. Therefore, to consider how local ecosystem processes are interlinked by dispersal of fungi, we need to take into account “metacommunity-scale” networks of plant-fungus associations [35]. Within a dataset of multiple local communities (e.g., [25]), fungal species that occur in multiple localities may interlink local networks of plant-fungus associations. Among them, some species that not only have broad geographic ranges but also are associated with diverse host plant species would be placed at the core positions of a metacommunity-scale network [35]. Such “metacommunity hub” fungi would be major drivers of the synchronization and restructuring of local ecosystem processes (*sensu* [50]), and hence their functional roles need to be investigated with priority [35]. Moreover, in the screening of mycorrhizal and endophytic fungi that can be used in agriculture and ecosystem restoration programs [17, 20, 51], analytical pipelines for identifying metacommunity hubs will help us explore species that are potentially applied (inoculated) to diverse plant species over broad geographic ranges of farmlands, forests, or grasslands. Nonetheless, despite the potential importance of metacommunity hubs in both basic and applied microbiology, few studies have examined metacommunity-level networks of plant-symbiont associations.

By compiling Illumina sequencing datasets of root-associated fungi [52], we herein inferred a metacommunity-level network of below-ground plant-fungus associations and thereby explored metacommunity hubs. Our metadata consisted of plant-fungus association data in eight forest localities across the entire range of the Japanese Archipelago, including 150 plant species/taxa and 8,080 fungal operational taxonomic units (OTUs) in temperate and subtropical regions. Based on the information of local- and metacommunity-level networks, each of the fungal OTUs was evaluated in light of its topological positions. We then examined whether fungal OTUs placed at the core of local-level plant-fungus networks could play key topological roles within the metacommunity-level network. Overall, this study uncover how diverse taxonomic groups of mycorrhizal and endophytic fungi can form metacommunity-scale networks of below-ground plant-fungus associations, providing a basis for analyzing complex spatial processes of species-rich host-microbe systems.

## Methods

### Terminology

While a single type of plant-fungus interactions is targeted in each of most mycological studies (e.g., arbuscular mycorrhizal symbiosis or ectomycorrhizal symbiosis), we herein analyze the metadata including multiple categories of below-ground plant-fungus associations [52]. Because arbuscular mycorrhizal, ectomycorrhizal, and endophytic fungi, for example, vary in their microscopic structure within plant tissue [38], it is impossible to develop a general criterion of mutualistic/antagonistic interactions for all those fungal functional groups. Therefore, we used the phrase “associations” instead of “interactions” throughout the manuscript when we discuss patterns detected based on the Illumina sequencing metadata of root-associated fungi. Consequently, our results represented not only mutualistic or antagonistic interactions but also neutral or commensalistic interactions [24, 53, 54]. Our aim in this study is to gain an overview of the metacommunity-scale plant-fungus associations, while the nature of respective plant-fungus associations should be evaluated in future inoculation experiments.

### Data

We compiled the Illumina (MiSeq) sequencing data collected in a previous study [52], in which community-scale statistical properties of below-ground plant-fungus associations were compared among eight forest localities (four cool-temperate, one warm-temperate, and three subtropical forests) across the entire range of the Japanese Archipelago (45.042-24.407 °N; Fig. 1). In each forest, 2-cm segment of terminal roots were sampled from 3-cm below the soil surface at 1-m horizontal intervals [52]. Those root samples were collected irrespective of their morphology and mycorrhizal type: hence, the samples as a whole represented below-ground relative abundance of plant species in each forest community. Based on the sequences of the genes encoding the large subunit of ribulose-1,5-bisphosphate carboxylase (*rbcL*) and the internal transcribed spacer 1 (ITS1) of the ribosomal RNA region, host plant species were identified, although there were plant root samples that could not be identified to species with the *rbcL* and ITS1 regions [52]. The sequencing data are available through DDBJ Sequence Read Archives (accession: DRA006339).

**Fig. 1.**
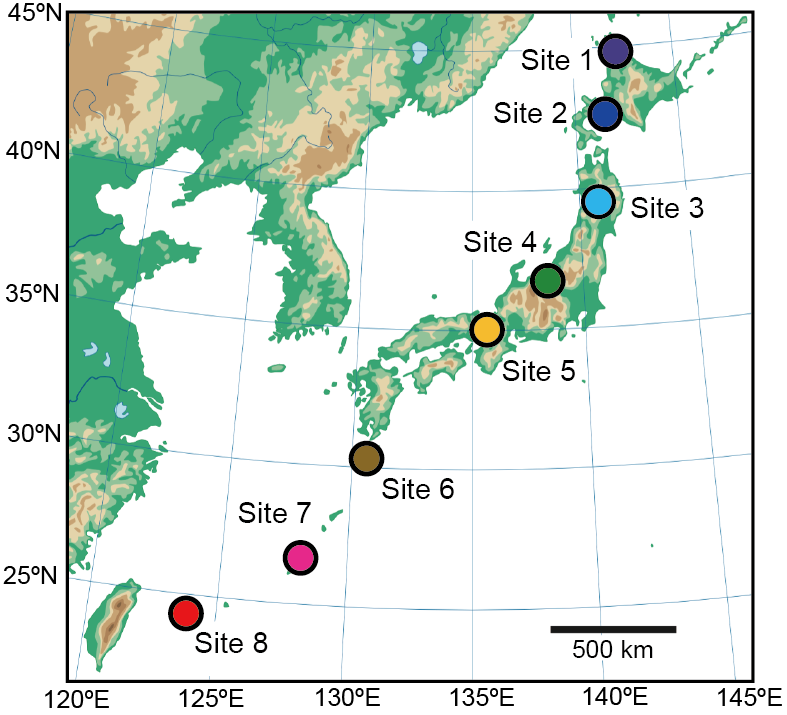
Study sites examined in this study. Across the entire range of the Japanese Archipelago, root samples were collected in four cool-temperate forests (sites 1-4), one warm-temperate forest (site 5), and three subtropical forests (sites 6-8).

The Illumina sequencing reads of the fungal ITS1 region were processed using the program Claidnet [55, 56] as detailed in the data-source study [52]: the Unix scripts used are available as Additional file 1. The primers used were designed to target not only Ascomycota and Basidiomycota but also diverse non-Dikarya (e.g., Glomeromycota) taxa [57]. In most studies analyzing community structure of Ascomycota and Basidiomycota fungi, OTUs of the ITS region are defined with a cut-off sequence similarity of 97% [22, 58, 59] (see also [60]). Meanwhile, Glomeromycota fungi generally have much higher intraspecific ITS-sequence variation than other taxonomic groups of fungi [61]. Consequently, we used 97% and 94% cut-off sequence similarities for defining non-Glomeromycota and Glomeromycota fungal OTUs, respectively [52]. The OTUs were then subjected to reference database search with the query-centric auto-*k*-nearest-neighbor algorithm [55, 56] and subsequent taxonomic assignment with the lowest common ancestor algorithm [62]. Based on the inferred taxonomy, the functional group of each fungal OTU was inferred using the program FUNGuild 1.0 [63].

After a series of bioinformatics and rarefaction procedures, 1,000 fungal ITS reads were obtained from each of the 240 samples collected in each forest locality (i.e., 1,000 reads × 240 samples × 8 sites). A sample (row) × fungal OTU (column) data matrix, in which a cell entry depicted the number of sequencing reads of an OTU in a sample, was obtained for each local forest (“sample-level” matrix) (Additional file 2: Data S2). Each local sample-level matrix was then converted into a “species-level” matrix, in which a cell entry represented the number of root samples from which associations of a plant species/taxa (row) and a fungal OTU (columns) was observed: 17-55 plant species/taxa and 1,149-1,797 fungal OTUs were detected from the local species-level matrices (Additional file 3: Data S3). In total, the matrices included 150 plant species/taxa and 8,080 fungal OTUs (Additional file 4: Data S4).

### Local networks

Among the eight forest localities, variation in the order-level taxonomic compositions were examined with the permutational analysis of variance (PERMANOVA; [64]) and the permutational analysis for the multivariate homogeneity of dispersions (PERMDISP; [65]) with the “adonis” and “betadisper” functions of the vegan 2.4-3 package [66] of R 3.4.1 [67], respectively. The *β*-diversity values used in the PERMANOVA and PERMDISP analyses were calculated with the “Bray-Curtis” metric based on the sample-level matrices (Additional file 2: Data S2). Note that the “Raup-Crick” *β*-diversity metric [68], which controls *α*-diversity in community data but requires computationally intensive randomization, was not applicable to our large metadata. Geographic variation in the compositions of fungal functional groups was also evaluated by PERMANOVA and PERMDISP analyses. The R scripts for the PERMANOVA and PERMDISP analyses are available as Additional file 5.

For each of the eight local forests, the network structure of below-ground plant-fungus associations was visualized based on the species-level matrix (Additional file 3: Data S3) using the program GePhi 0.9.1 [69] with the “ForceAtlas2” layout algorithm [70]. Within the networks, the order-level taxonomy of fungal OTUs was highlighted.

To evaluate host ranges of each fungal OTU in each local forest, we first calculated the *d’* metric of interaction specificity [71]. However, estimates of the *d’* metric varied considerably among fungal OTUs observed from small numbers of root samples (Additional file 6; Figure S1) presumably due to overestimation or underestimation of host preferences for those rare OTUs. Therefore, we scored each fungal OTU based on their topological positions within each local network by calculating network centrality indices (degree, closeness, betweenness, and eigenvector centralities metrics of network centrality; [31]). Among the centrality metrics, betweenness centrality, which measures the extent to which a given nodes (species) is located within the shortest paths connecting pairs of other nodes in a network [72], is often used to explore organisms with broad host or partner ranges [35]. Thus, in each local network, fungal OTUs were ranked based on their betweenness centrality scores (local betweenness).

### Metacommunity-scale network

By compiling the species-level matrices of the eight local forests, the topology of the metacommunity-scale network of plant-fungus associations was inferred. In general, species interaction (association) networks of local communities can be interconnected by species that appear in two or more local networks, thereby merged into a metacommunity-scale network [35]. In our data across the eight local forests, 2,109 OTUs out of the 8,080 fungal OTUs appeared in two or more localities. Therefore, we could infer the topology of a metacommunity-scale network, in which the eight local networks were combined by the 2,109 fungal OTUs. In the metacommunity-scale network, plant species/taxa observed in different localities were treated as different network nodes because our purpose in this study was to explore fungi that potentially play key roles in synchronizing local ecosystem processes [35]. In total, 227 plant nodes representing local populations of 150 plant species/taxa were included in the metacommunity-scale network.

We then screened for fungal OTUs with broad geographic and host ranges based on the betweenness centrality scores of respective fungal OTUs within the metacommunity network (metacommunity betweenness, *B*_meta_). In general, species with highest metacommunity betweenness scores not only occur in local communities over broad biotic/abiotic environmental conditions but also are associated with broad ranges of host/partner species [35]. Possible relationship between local- and metacommunity-scale topological roles was then examined by plotting local and metacommunity betweenness scores ( *B*_local_ and *B*_meta_) of each fungal OTUs on a two-dimensional surface. To make the betweenness scores vary from 0 to 1, betweenness centrality of a fungal OTU *i* was standardized in each of the local- and metacommunity-scale networks as follows:

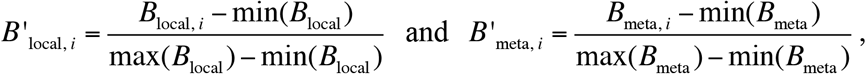

where *B*_local, *i*_ and *B*_meta, *i*_ were raw estimates of local- and metacommunity-scale betweenness of a fungal OTU *i*, and min() and max() indicated minimum and maximum values, respectively. For local betweenness of each OTU, a mean value across local networks was subsequently calculated 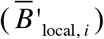: the local communities from which a target OTU was absent was omitted in the calculation of mean local betweenness. On the two-dimensional surface, the OTUs were then classified into four categories: metacommunity hubs having high betweenness in both local- and metacommunity-scale networks (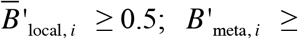 0.5), metacommunity connectors that had broad geographic ranges but displayed low local betweenness 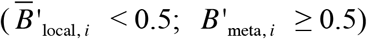, local hubs that had high betweenness in local networks but not in the metacommunity-scale network 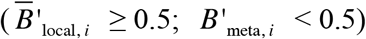, and peripherals with low betweenness at both local and metacommunity levels 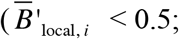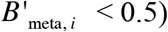 [35]. Approximately, 1-2% of fungal OTUs show betweenness scores higher than 0.5 in each local or metacommunity network, while the threshold value can be changed depending on the purpose of each study [35].

In addition to metacommunity hubs within the metacommunity-scale network representing all the eight localities, those within the metacommunity-scale network representing northern (sites 1-4) or southern (sites 5-8) four localities were also explored. This additional analysis allowed us to screen for fungal OTUs that potentially adapted to broad ranges of biotic and abiotic environments within northern (cool-temperate) or southern (warm-temperate or subtropical) part of Japan.

## Results

### Local networks

Among the eight forest localities, order-level taxonomic compositions of fungi varied significantly (PERMANOVA; *F*_model_ = 35.7, *P* < 0.001), while the differentiation of community structure was attributed at least partly to geographic variation in among-sample dispersion (PERMDISP; *F* = 13.2, *P* < 0.001) (Fig. 2a). Compositions of fungal functional groups were also differentiated among the eight localities (PERMANOVA; *F*_model_ = 34.9, *P* < 0.001), while within-site dispersion was significantly varied geographically (PERMDISP; *F* = 9.2, *P* < 0.001) (Fig. 2b). The proportion of ectomycorrhizal fungal orders, such as Russulales, Thelephorales, and Sebacinales, was higher in temperate forests than in subtropical forests, while that of arbuscular mycorrhizal fungi increased in subtropical localities (Fig. 2). The proportion of the ascomycete order Helotiales, which has been known to include not only ectomycorrhizal but also endophytic, saprotrophic, and ericoid mycorrhizal fungi [73], was higher in northern localities. In contrast, Diaporthales, which has been considered as predominantly plant pathogenic taxon [74] (but see [75]), was common in subtropical forests but not in others.

**Fig. 2.**
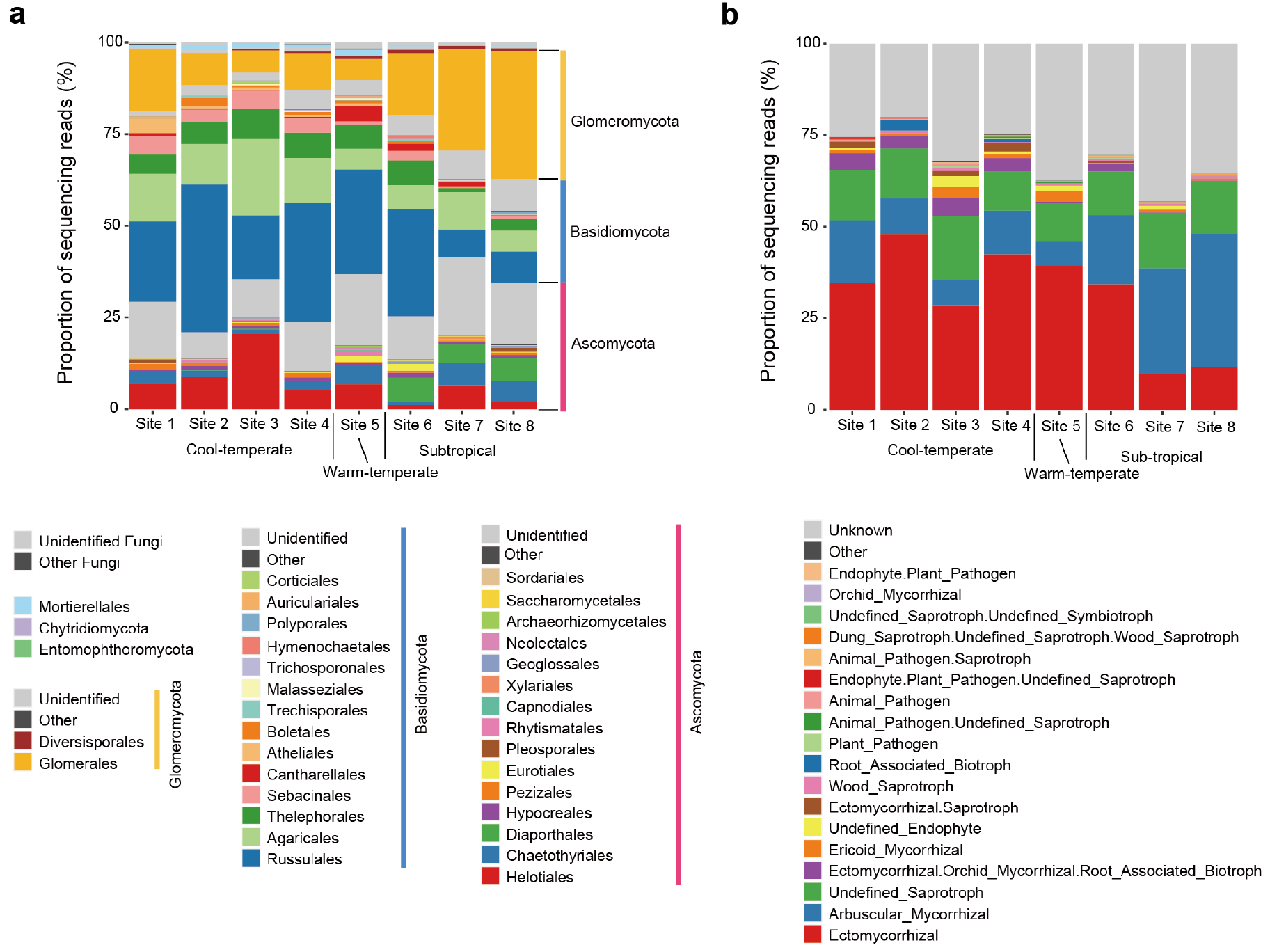
Compositions of fungal taxa and functional groups in each forest. **a** Order-level taxonomic composition of fungal OTUs in each locality. The number of fungal OTUs detected is shown in a parenthesis for each forest. **b** Functional-group composition. The fungal functional groups were inferred by the program FUNGuild [63].

In each of the eight local networks depicting plant-fungus associations, some fungal OTUs were located at the central positions of the network, while others are distributed at peripheral positions (Additional file 7; Figure S2). Specifically, fungal OTUs belonging to the ascomycete orders Chaetothyriales (e.g., *Cladophialophora* and *Exophiala*) and Helotiales (e.g., *Rhizodermea*, *Pezicula*, *Rhizoscyphus*, and *Leptodontidium*) as well as some *Mortierella* OTUs had high betweenness centrality scores in each of the cool-temperate forests (Fig. 3a-b). In contrast, arbuscular mycorrhizal fungi (Glomeromycota) were common among OTUs with highest betweenness scores in subtropical forests (Fig. 3a-c). Some fungi in the ascomycete order Hypocreales (e.g., *Trichoderma*, *Ilyonectria*, *Simplicillium*, and *Calonectria*) also had high betweenness scores in some temperate and subtropical forests (Fig. 3b).

**Fig. 3.**
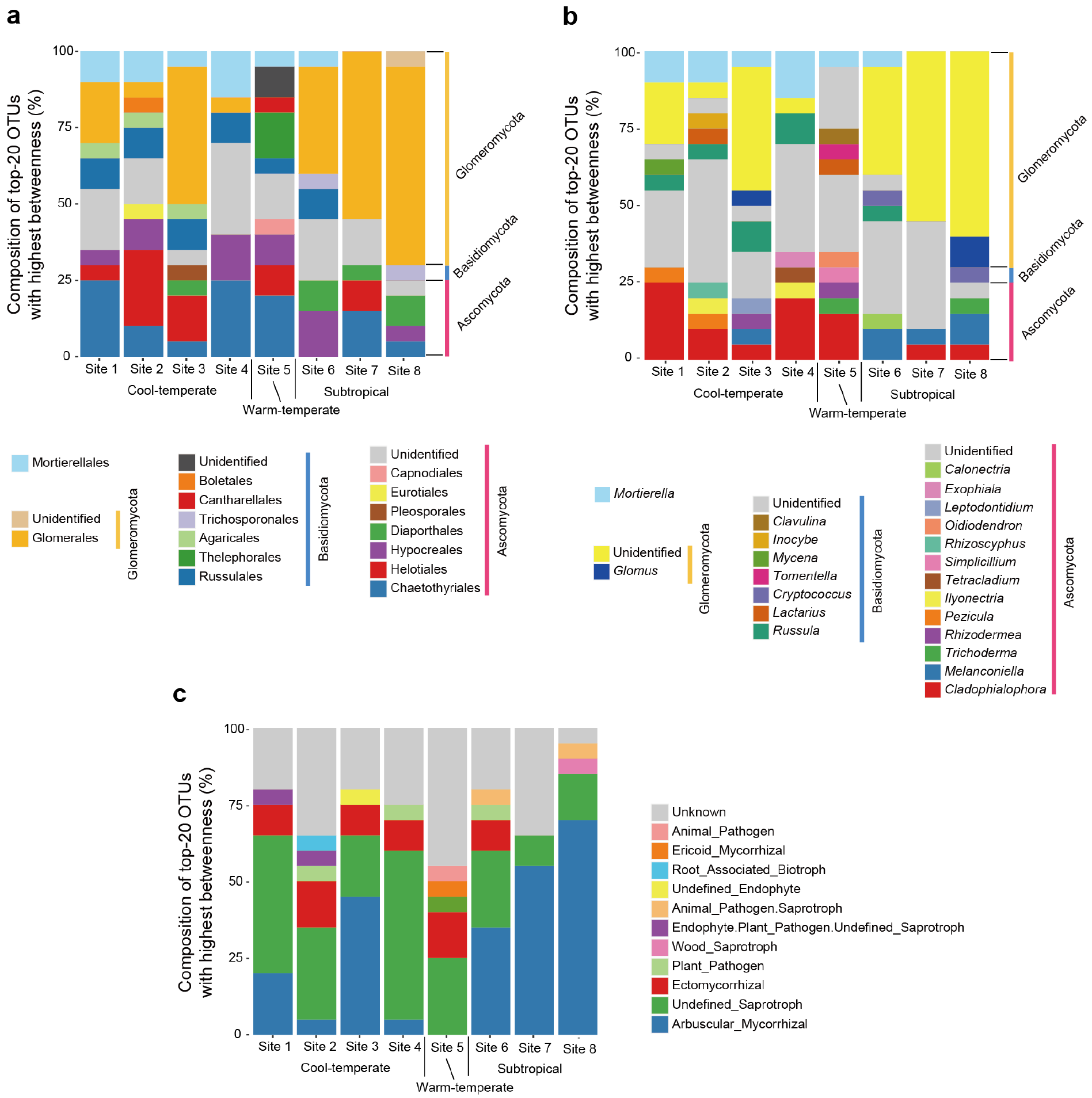
Fungal OTUs with highest local betweenness. **a** Order-level taxonomic composition of top-20 OTUs with highest local betweenness in each forest. See Data S4 (Additional file 4) for betweenness scores of all fungal OTUs in respective local forests. **b** Genus-level taxonomic composition of top-20 OTUs with highest local betweenness. **c** Functional-group composition of top-20 OTUs with highest local betweenness.

### Metacommunity-scale network

In the metacommunity-scale network representing the connections among the eight local networks, not only arbuscular mycorrhizal but also saprotrophic/endophytic fungi were placed at the central topological positions (Fig. 4; Additional file 8; Figure S3). Among non-Glomeromycota OTUs, *Mortierella* (Mortierellales), *Cryptococcus* (Trichosporonales; the Blast top-hit fungus in the NCBI database was recently moved to *Saitozyma* (Tremellales); [76]), *Malassezia* (Malasseziales), *Oidiodendron* (incertae sedis), *Trichoderma* (Hypocreales), and a fungus distantly allied to *Melanconiella* (Diaporthales) displayed highest metacommunity betweenness (Table 1). Among the OTUs with high metacommunity betweenness, only a *Mortierella* OTU was designated as a metacommunity hub (i.e., 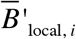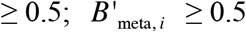 and others had low betweenness scores at the local community level (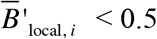; Fig. 5a).

**Fig. 4.**
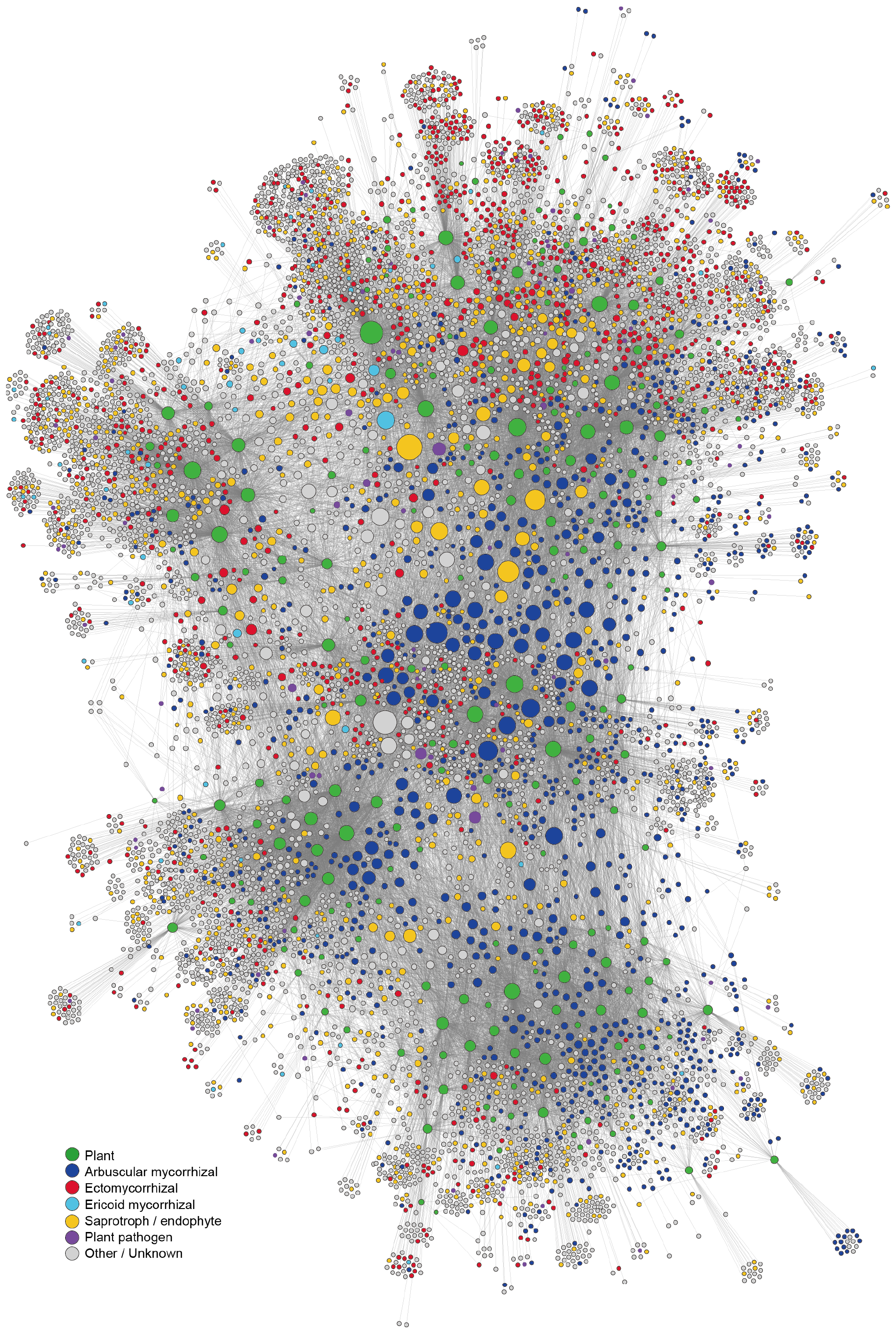
Metacommunity-scale network including all the eight local forests. The size of circles roughly represents relative scores of betweenness centrality. The functional groups of fungi inferred with the program FUNGuild [63] were organized into six categories: i.e., arbuscular mycorrhizal (bue), ectomycorrhizal (red), ericoid mycorrhizal (skyblue), saprotrophic/endophytic (yellow), plant pathogenic (purple), and other/unknown fungi (grey) (Additional file 4; Data S4). For plant species/taxa (green), the geographic information of source populations is indicated in Additional file 8 (Figure S3).

**Table 1.** Top-10 list of non-Glomeromycota OTUs with highest betweenness within the metacommunity networks. In each of the three metacommunity-scale networks examined (full, cool-temperate, and warm-temperate/subtropical), fungal OTUs were ranked based on their betweenness centrality scores. As taxonomic information of Glomeromycota OTUs with high betweenness scores was redundant (e.g., *Glomus* spp. or Glomeraceae spp.), the top-10 list of non-Glomeromycota OTUs is shown. Taxonomy information of each OTU was inferred based on the query-centric auto-*k*-nearest-neighbor algorithm of reference database search [55, 56] and subsequent taxonomic assignment with the lowest common ancestor algorithm [62]. The results of the NCBI nucleotide Blast are also shown. For simplicity, the functional groups of fungi inferred with the program FUNGuild [63] were organized into several categories. See Data S4 (Additional file 4) for details of the categories and for full results including Glomeromycota and other fungal OTUs.

| OTU | Phylum | Class | Order | Family | Genus | Category | NCBI Blast top hit | Accession | Cover | Identity |
| --- | --- | --- | --- | --- | --- | --- | --- | --- | --- | --- |
| Full (8sites) |  |  |  |  |  |  |  |  |  |  |
| F_0042* | - | - | Mortierellales | Mortierellaceae | *Mortierella* | Saprotroph/Endophyte | *Mortierella humilis* | KP714537 | 100% | 100% |
| F_0381 | Basidiomycota | Tremellomycetes | Trichosporonales | Trichosporonaceae | *Cryptococcus* | Others_Unknown | *Saitozyma podzolica*† | KY320605 | 92% | 99% |
| F_0079 | Ascomycota | Sordariomycetes | Hypocreales | Nectriaceae | - | Saprotroph/Endophyte | *Ilyonectria protearum* | NR_152890 | 99% | 100% |
| F_0489 | - | - | Mortierellales | Mortierellaceae | *Mortierella* | Saprotroph/Endophyte | *Mortierella* sp. | KM113754 | 100% | 100% |
| F_0010 | Ascomycota | Leotiomycetes | - | Myxotrichaceae | *Oidiodendron* | Ericoid_Mycorrhizal | *Oidiodendron maius* | LC206669 | 100% | 100% |
| F_0368 | Basidiomycota | Malasseziomycetes | Malasseziales | Malasseziaceae | *Malassezia* | Others_Unknown | *Malassezia restricta* | KT809059 | 100% | 100% |
| F_0623 | - | - | Mortierellales | Mortierellaceae | *Mortierella* | Saprotroph/Endophyte | *Mortierella gamsii* | KY305027 | 100% | 100% |
| F_1188 | Basidiomycota | Tremellomycetes | Trichosporonales | Trichosporonaceae | *Cryptococcus* | Others_Unknown | *Saitozyma podzolica*† | KY320605 | 92% | 99% |
| F_0007 | Ascomycota | Sordariomycetes | Diaporthales | Melanconidaceae | *Melanconiella* | Saprotroph/Endophyte | *Melanconiella elegans* | KJ173701 | 100% | 85% |
| F_0485 | Ascomycota | Sordariomycetes | Hypocreales | Hypocreaceae | *Trichoderma* | Saprotroph/Endophyte | *Trichoderma* sp. | HG008760 | 100% | 100% |

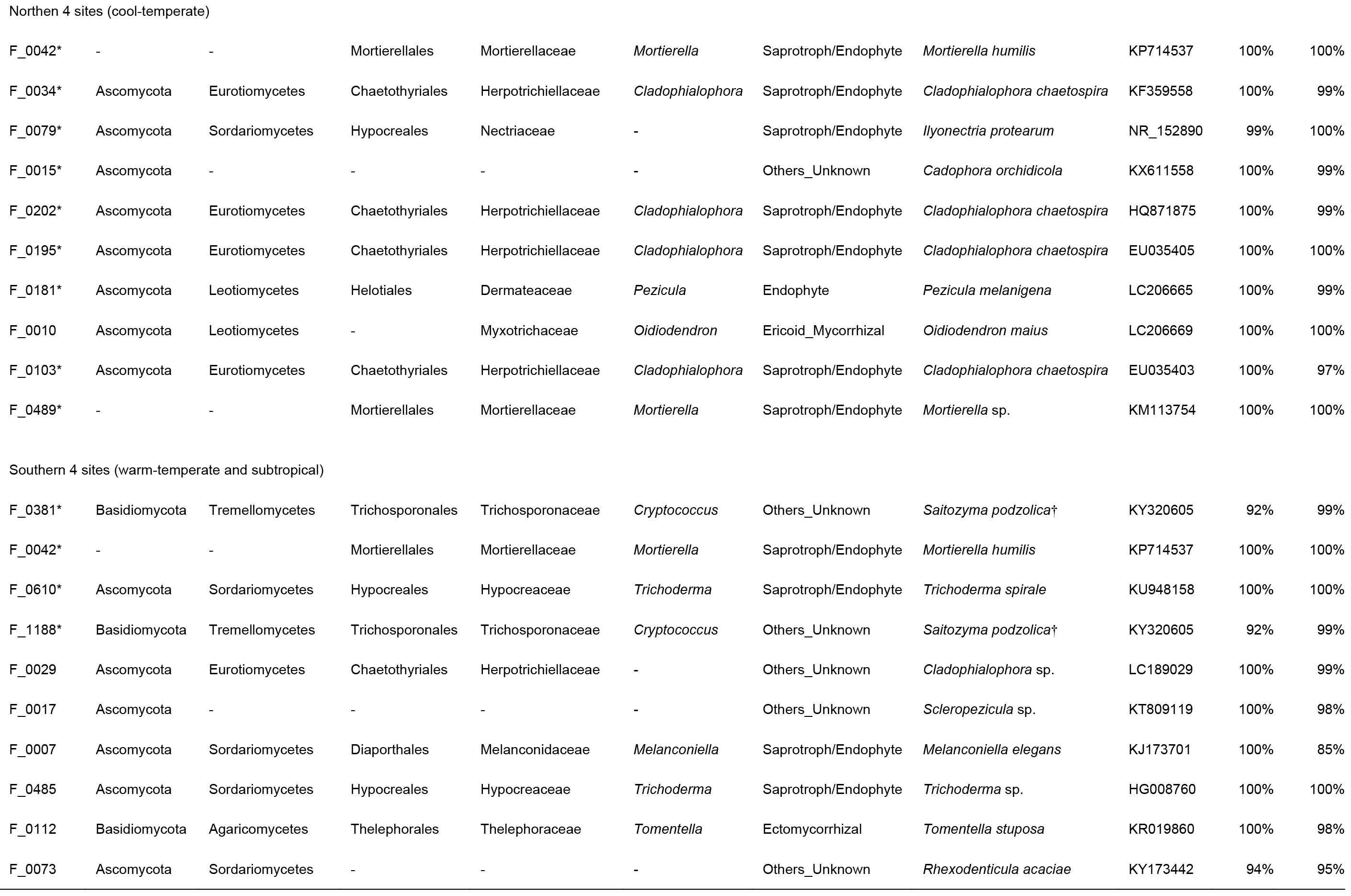

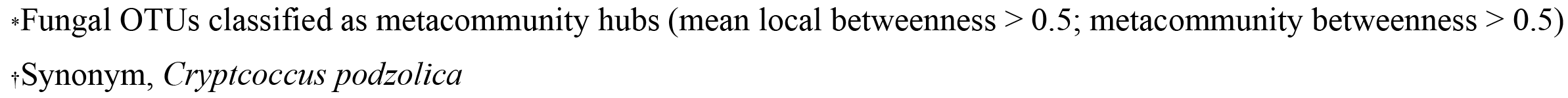

In the metacommunity-scale network representing the four cool-temperate forests (sites 1-4), many saprotrophic/endophytic fungal OTUs were associated with diverse plant species/taxa, located at the central topological positions within the network topology (Additional file 9; Figure S4; Fig. 5b). The list of these fungi with high metacommunity betweenness involved OTUs in the genera *Mortierella*, *Cladophialophora* (Chaetothyriales), *Pezicula* (Helotiales), and *Oidiodendron* as well as OTUs allied to *Ilyonectria protearum* (Nectriales) and *Cadophora orchidicola* (Helotiales) (Table 1). Most of those fungal OTUs also had high metacommunity betweenness, designated as metacommunity hubs (Fig. 5b).

**Fig. 5.**
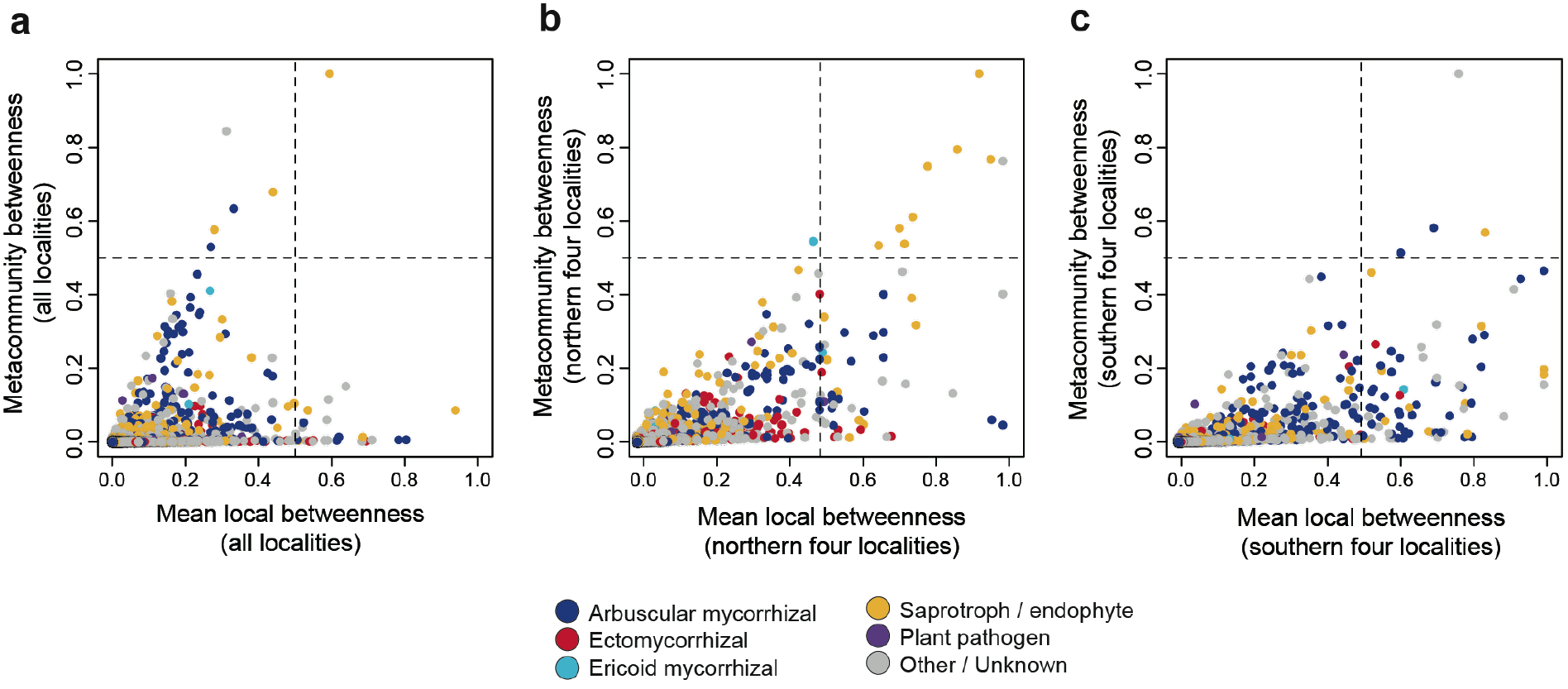
Relationship between local- and metacommunity-level betweenness. **a** Full meatcommunity. On the horizontal axis, the mean values of betweenness centrality scores across all the eight local forests are shown for respective fungal OTUs. On the vertical axis, the betweenness scores within the metacommunity-scale network consisting of the eight localities (Fig. 4) are shown for respective OTUs. **b** Metacommunity of cool-temperate forests. For the sub-dataset consisting of the four cool-temperate forests (Additional file 9: Figure S4), mean local betweenness and metacommunity betweenness are shown on the horizontal and vertical axes, respectively. **c** Metacommunity of warm-temperate and subtropical forests. For the sub-dataset consisting of the warm-temperate forest and the three subtropical forests (Additional file 10: Figure S5), mean local betweenness and metacommunity betweenness are shown on the horizontal and vertical axes, respectively.

In the metacommunity-scale network consisting of the warm-temperate and subtropical forests (sites 5-8), arbuscular mycorrhizal and saprotrophic/endophytic fungi were placed at the hub positions (Additional file 10; Figure S5; Fig. 5c). The list of non-Glomeromycota OTUs with highest metacommunity betweenness included *Saitozyma* (*Cryptococcus*), *Mortierella*, *Trichoderma*, and *Tomentella* as well as OTUs allied to *Cladophialophora*, *Scleropezicula* (Helotiales), *Melanconiella* (Diaporthales), and *Rhexodenticula* (incertae sedis) (Table 1). Among the taxa, *Saitozyma* and *Mortierella* included OTUs classified as metacommunity hubs (Fig. 5c; Table 1). In an additional analysis of a metacommunity-scale network including only the three subtropical forests (sites 6-8), similar sets of fungal taxa were highlighted (Additional file 11; Table S1). The detailed information of the network index scores examined in this study is provided in Data S3 (Additional file 4: Data S4).

## Discussion

Based on the metadata of root-associated fungi across the Japanese Archipelago, we herein inferred the structure of a network representing metacommunity-scale associations of 150 plant species/taxa and 8,080 fungal OTUs. Our analysis targeted diverse functional groups of fungi such as arbuscular mycorrhizal, ectomycorrhizal, ericoid-mycorrhizal, saprotrophic/endophytic, and pathogenic fungi, which have been analyzed separately in most previous studies on plant-fungus networks. The comprehensive analysis of below-ground plant-fungus associations allowed us to explore metacommunity hub fungi, which not only occurred over broad geographic ranges but also had broad host ranges in respective local communities. Consequently, this study highlights several taxonomic groups of fungi potentially playing key roles in synchronizing metacommunity-scale processes of temperate and/or subtropical forests.

In the metacommunity-scale network representing all the eight local forests (Fig. 4), fungi in several saprotrophic or endophytic taxa showed higher betweenness centrality scores than other fungi (Table 1). *Mortierella* is generally considered as a saprotrophic lineage [77] but it also includes fungi contributing to the growth and pathogen resistance of plants [78–80]. A phosphate solubilizing strain of *Mortierella*, for example, increases shoot and root growth of host plants under salt stress, especially when co-inoculated with an arbuscular mycorrhizal fungus [78]. In addition, polyunsaturated fatty acids produced by some *Mortierella* species are known to increase resistance of plants against phytopathogens [79, 80]. Fungi in the genus *Trichoderma* are commonly detected and isolated from the rhizosphere [77, 81]. Many of them inhibit the growth of other fungi, often used in the biological control of phytopathogens [82–84]. Some of them are also reported to suppress root-knot nematodes [85] or to promote root growth [86]. The analysis also highlighted basidiomycete yeasts in the genus *Saitozyma* or *Cryptococcus* (teleomorph = *Filobasidiella*), which are often isolated from soil [22, 87] as well as both above-ground and below-ground parts of plants [88–91].

Along with those possibly saprotrophic or endophytic taxa, ericoid mycorrhizal and phytopathogenic taxa of fungi displayed relatively high betweenness scores within the metacommunity-scale network representing all the eight local forests (Table 1). Specifically, *Oidiodendron* (teleomorph = *Myxotrichum*) is a taxon represented by possibly ericoid mycorrhizal species (*O. maius* and *O. griseum*) [92, 93], although fungi in the genus are found also from roots of non-ericaceous plants and soil [94]. On the other hand, fungi in the family Nectriaceae are known to cause black foot disease [95], often having serious damage on economically important woody plants [96, 97]. Although we collected seemingly benign roots in the study forests, some samples may be damaged by those pathogens. Alternatively, some lineages of Nectriaceae fungi may be associated with plant hosts non-symptomatically, having adverse effects context-dependently.

Although these fungi were candidates of metacommunity hubs, which are characterized by broad geographic ranges and host plant ranges, none except but a *Mortierella* OTU had high betweenness scores at both local and metacommunity levels (Fig. 5a). This result suggests that even if some fungi have broad geographic ranges across the Japanese Archipelago, few played important topological roles in each of the local networks representing plant-fungus associations. In other words, fungi that can adapt to biotic and abiotic environments in forest ecosystems throughout cool-temperate, warm-temperate, and subtropical regions are rare.

Therefore, we also explored fungi with broad geographic and host ranges within the metacommunities representing northern (cool-temperate) and southern (warm-temperate and subtropical) regions of Japan. In the metacommunity consisting of the four cool-temperate forests (Additional file 9; Figure S4), fungal OTUs in the genera *Mortierella*, *Cladophialophora*, and *Pezicula* as well as those allied to *Ilyonectria* and *Cadophora* had highest betweenness at both local and metacommunity levels, classified as metacommunity hubs (Fig. 5b; Table 1). Among them, *Cladophialophora* is of particular interest because it has been known as a lineage of “dark septate endophytes” [98–100] (*sensu* [14, 15, 101]). A species within the genus, *C. chaetospira* (= *Heteroconium chaetospira*), to which high-betweenness OTUs in our data were closely allied, has been known not only to provide nitrogen to host plants but also to suppress pathogens [12, 16, 102]. Likewise, the Helotiales genus *Pezicula* (anamorph = *Cryptosporiopsis*) includes endophytic fungi [103–105], some of which produce secondary metabolites suppressing other microbes in the rhizosphere [106, 107]. Our finding that some of *Cladophialophora* and *Pezicula* fungi could be associated with various taxonomic groups of plants over broad geographic ranges highlights potentially important physiological and ecological roles of those endophytes at the community and metacommunity levels.

In the southern metacommunity networks consisting of warm-temperate and subtropical forests (Additional file 10; Figure S5), some arbuscular mycorrhizal OTUs and *Saitozyma* (*Cryptococcus*) and *Mortierella* OTUs had high betweenness scores at both local and metacommunity levels, designated as metacommunity hubs (Fig. 5c; Table 1). Given the above-mentioned prevalence of fungal OTUs allied to *Cladophialophora chaetospira* in the cool-temperate metacommunity, the contrasting list of metacommunity hubs in the southern (warm-temperate-subtropical) metacommunity implies that different taxonomic and functional groups of fungi play major metacommunity-scale roles in different climatic regions. This working hypothesis is partially supported by previous studies indicating endemism and vicariance in the biogeography of fungi and bacteria [108, 109], promoting conceptual advances beyond the classic belief that every microbe is everywhere but the environment selects microbes colonizing respective local communities [110].

The roles of those metacommunity hubs detected in this study are of particular interest from the aspect of theoretical ecology. Hub species connected to many other species in an ecosystem often integrate “energy channels” [111] within species interaction networks, having great impacts on biodiversity and productivity of the ecosystems [35]. The concept of “keystone” or “foundation” species [112, 113] can be extended to the metacommunity level, thereby promoting studies exploring species that restructure and synchronize ecological (and evolutionary) dynamics over broad geographic ranges [35]. Given that below-ground plant-fungus symbioses are key components of the terrestrial biosphere [1, 2], identifying fungal species that potentially have great impacts on the metacommunity-scale processes of such below-ground interactions will provide crucial insights into the conservation and restoration of forests and grasslands. We here showed that the list of metacommunity hubs could involve various lineages of endophytic fungi, whose ecosystem-scale functions have been underappreciated compared to those of mycorrhizal fungi. As those endophytic fungi are potentially used as inoculants when we reintroduce plant seedlings in ecosystem restoration programs [20, 51], exploring fungi with highest potentials in each climatic/biogeographic region will be a promising direction of research in conservation biology.

The finding that compositions of metacommunity hubs could vary depending on climatic regions also gives key implications for the application of endophytes in agriculture. Although a number of studies have tried to use endophytic fungi and/or bacteria as microbial inoculants in agriculture [17, 18, 114], such microbes introduced to agroecosystems are often outcompeted and replaced by indigenous (resident) microbes [115, 116]. Moreover, even if an endophytic species or strain increases plant growth in pot experiments under controlled environmental conditions, its effects in the field often vary considerably depending on biotic and abiotic contexts of local agroecosystems [17] (see also [117]). Therefore, in the screening of endophytes that can be used in broad ranges of biotic and abiotic environmental conditions, the metacommunity-scale network analysis outlined in this study will help us find promising candidates out of thousands or tens of thousands microbial species in the wild. Consequently, to find promising microbes whose inocula can persist in agroecosystems for long time periods, exploration of metacommunity hubs needs to be performed in respective climatic or biogeographic regions.

For more advanced applications in conservation biology and agriculture, continual improvements of methods for analyzing metacommunity-scale networks are necessary. First, while the fungal OTUs in our network analysis was defined based on the cut-off sequence similarities used in other studies targeting “species-level” diversity of fungi [59, 61], physiological functions can vary greatly within fungal species or species groups [14, 118].
Given that bioinformatic tools that potentially help us detect single-nucleotide-level variation are becoming available [119], the resolution of network analyses may be greatly improved in the near future. Second, although some computer programs allow us to infer functions of respective microbial OTUs within network data [63, 120], the database information of microbial functions remains scarce. To increase the coverage and accuracy of automatic annotations of microbial functions, studies describing the physiology, ecology, and genomes of microbes should be accelerated. With improved reference databases, more insights into the metacommunity-scale organization of plant-fungus associations will be obtained by reanalyzing the network data by compiling enhanced information of fungal functional groups. Third, as the diversity and compositions of plant-fungus associations included in a network can depend on how we process raw samples, special care is required in the selection of methods for washing and preparing root (or soil) samples. By sterilizing root samples with NaClO [121], for example, we may be able to exclude fungi or bacteria that are merely adhering to root surfaces. Meanwhile, some of those fungi and bacteria on root surfaces may play pivotal physiological roles in the growth and survival of plants [122]. Accordingly, it would be productive to compare network topologies of plant-microbe associations among different source materials by partitioning endosphere, rhizoplane, and rhizosphere microbial samples with a series of sample cleaning processes using ultrasonic devices [123]. Fourth, although this study targeted fungi associated with roots, our methods can be easily extended to network analyses involving other groups of microbes. By simultaneously analyzing the prokaryote 16S rRNA region [123–125] with the fungal ITS region, we can examine how bacteria, archaea, and fungi are involved in below-ground webs of symbioses. Fifth, not only plant-microbe associations but also microbe-microbe interactions can be estimated with network analytical frameworks. Various statistical pipelines have been proposed to infer how microbes interact with each other in facilitative or competitive ways within host macroorganisms [37, 126, 127]. Overall, those directions of analytical extensions will enhance our understanding of plant microbiome dynamics in nature.

## Conclusions

By compiling datasets of below-ground plant-fungus associations in temperate and subtropical forest ecosystems, we explored metacommunity-hub fungi, which were characterized by broad geographic and host ranges. Such metacommunity-scale analyses are expected to provide bird’s-eye views of complex plant-microbe associations, highlighting plant-growth-promoting microbes that can be applied to diverse plant taxa in various environments. Given that endophytic fungi promoting the growth and pathogen resistance of host plants can be isolated from forest soil (e.g., *Cladophialophora chaetospira* [99]), the list of metacommunity-hub endophytic fungi featured in this study itself may include prospective species to be used in agriculture. By extending the targets of such network analyses to diverse types of plant-associated microbes (e.g., phyllosphere fungi and bacteria [75, 124, 128]) in various climatic/biogeographic regions, a solid basis for managing plant microbiomes will be developed.

## Abbreviations

DDBJ: DNA Data Bank of Japan
ITS: internal transcribed spacer
OTU: Operational taxonomic unit
PERMANOVA: permutational analysis of variance
PERMDISP: permutational analysis for the multivariate homogeneity of dispersions
rRNA: ribosomal ribonucleic acid.

## Acknowledgements

We thank Teshio Experimental Forest (Hokkaido University), Tomakomai Experimental Forest (Hokkaido University), Sugadaira Research Station (Tsukuba University), Yona Field (Ryukyu University), Tropical Biosphere Research Center (Ryukyu University), and Forestry Agency of Japan for the permission of fieldwork.

## Availability of data and materials

The Illumina sequencing data were deposited to DNA Data Bank of Japan (DDBJ Sequence Read Archive: DRA006339). The raw data of fungal community structure and the fungal community matrices analyzed are available with the source study [52] and Additional files 2-4, respectively. The Unix scripts for the bioinformatic pipeline and the R scripts for the PERMANOVA and PERMDISP analyses are available as Additional files 1 and 4, respectively.

## Authors’ contributions

HT designed the work. HT, AST, and HS conducted fieldwork. HT performed the molecular experiments. HT wrote the manuscript with AST and HS.

## Consent for publication

Not applicable

## Ethics approval and consent to participate

Not applicable

## Additional files

**Additional file 1: Data S1.** Unix scripts for the bioinformatic pipeline.

**Additional file 2: Data S2.** Sample-level matrices of the eight forests examined.

**Additional file 3: Data S3.** Species-level matrices of plant-fungus associations.

**Additional file 4: Data S4.** Information of 8080 fungal OTUs analyzed.

**Additional file 5: Data S5.** R scripts for the PERMANOVA and PERMDISP analyses.

**Additional file 6: Figure S1.** Number of sequencing reads, interaction specificity, and local betweenness.

**Additional file 7: Figure S2.** Structure of plant-fungus networks in each local forest.

**Additional file 8 Figure S3.** Locality information within the full metacommunity-scale network.

**Additional file 9: Figure S4.** Metacommunity-scale network of cool-temperate forests.

**Additional file 10: Figure S5.** Metacommunity-scale network of warm-temperate and subtropical forests.

**Additional file 11: Table S1.** Top-10 list of non-Glomeromycota OTUs with highest betweenness within the subtropical metacommunity network.

